# Differential brain activity in visuo-perceptual regions during landmark-based navigation in young and healthy older adults

**DOI:** 10.1101/2020.03.13.990572

**Authors:** Stephen Ramanoël, Marion Durteste, Marcia Bécu, Christophe Habas, Angelo Arleo

**Author notes:** Co-first authorship (Stephen Ramanoël, Marion Durteste). **CA, Corresponding Author** Dr. Stephen Ramanoël, mail.

## Abstract

Older adults exhibit prominent impairments in their capacity to navigate, reorient in unfamiliar environments or update their path when faced with obstacles. This decline in navigational capabilities has traditionally been ascribed to memory impairments and dysexecutive function whereas the impact of visual aging has often been overlooked. The ability to perceive visuo-spatial information such as salient landmarks is essential to navigate in space efficiently. To date, the functional and neurobiological factors underpinning landmark processing in aging remain insufficiently characterized. To address this issue, this study used functional magnetic resonance imaging (fMRI) to investigate the brain activity associated with landmark-based navigation in young and healthy older participants. Twenty-five young adults (μ=25.4 years, σ=4.7; 7F) and twenty-one older adults (μ=73.0 years, σ=3.9; 10F) performed a virtual navigation task in the scanner in which they could only orient using salient landmarks. The underlying whole-brain patterns of activity as well as the functional roles of scene-selective regions, the parahippocampal place area (PPA), the occipital place area (OPA), and the retrosplenial cortex (RSC) were analyzed. We found that older adults’ navigational abilities were diminished compared to young adults’ and that the two age groups relied on distinct navigational strategies to solve the task. Better performance during landmark-based navigation was found to be associated with increased neural activity in an extended neural network comprising several cortical and cerebellar regions. Direct comparisons between age groups further revealed that young participants had enhanced anterior temporal activity. In addition, young adults only were found to recruit occipital areas corresponding to the cortical projection of the central visual field during landmark-based navigation. The region-of-interest analysis revealed increased OPA activation in older adult participants. There were no significant between-group differences in PPA and RSC activations. These results hint at the possibility that aging diminishes fine-grained information processing in occipital and temporal regions thus hindering the capacity to use landmarks adequately for navigation. This work helps towards a better comprehension of the neural dynamics subtending landmark-based navigation and it provides new insights on the impact of age-related visuo-spatial processing changes on navigation capabilities.

## 1. Introduction

Demographically, the 21st century is characterized by an unprecedented increase in the number of older adults within the worldwide population. There were 703 million people aged 65 years or over in 2019 and this number is projected to more than double by 2050 (United Nations, 2019). In parallel, we can expect a significant rise in the prevalence of neurodegenerative diseases such as Alzheimer’s and Parkinson’s diseases in the older population. In order to identify appropriate biomarkers, it is of critical importance that we gain a better understanding of brain changes in healthy aging. In this context, spatial navigation as a complex behavior encompassing perceptual and cognitive processes provides an ideal framework for the study of normal and pathological aging (Gazova et al., 2012; Lithfous et al., 2013; Allison et al., 2016; Laczo et al., 2017, 2018; Coughlan et al., 2018).

An extensive body of literature has highlighted a robust age-related decline in navigation ability in various species including rodents and non-human and human primates (Foster et al., 2012; Lester et al., 2017). Healthy older adults exhibit prominent impairments in their capacity to navigate efficiently, reorient or update their wayfinding behavior when faced with obstacles (Iaria et al., 2009; Moffat, 2009; Harris et al., 2012; Merhav et al., 2019). In real-world settings they are impaired at rapidly acquiring information about their surroundings leading to slower and more error-prone navigation than young adults (Kirasic, 1991; Wilkniss et al., 1997). In virtual reality (VR) paradigms older adults also choose inefficient routes, underestimate distances and make frequent turning errors (Adamo et al., 2012). Studies in VR have shed light on an age-related shift in the use of navigation strategies: older adults favor response over place-based strategies (Rodgers et al., 2012). A place-based strategy involves the formation of mental map-like representations of the absolute spatial position of the goal in relation to various stimuli within the environment. A response-based strategy refers to the process whereby an association between a specific stimulus and the goal location is formed. The choice of a navigation strategy critically depends on the visual information present in the environment (Foo et al., 2005; Ratliff and Newcombe, 2008). Indeed, successful navigation requires the perception and the integration of relevant spatial visual cues such as buildings or monuments, and the binding of these salient elements to directional information (Ekstrom, 2015; Epstein et al., 2017; Julian et al., 2018).

Spatial visual cues can be salient objects used as navigational landmarks or characteristics pertaining to the geometric shape of a space (Lester et al., 2017; Bécu et al., 2020). Landmarks can be conceptualized as discrete objects that are independent of the environment’s layout, such as a tree or a monument (Epstein and Vass, 2014). Landmarks’ size, stability and proximity to the goal are among the key factors that influence their use for navigation (Stankiewicz and Kalia, 2007; Auger et al., 2012; Auger and Maguire, 2018). Geometric cues encompass all the elements that are intrinsic to and continuous with the external limits of a space; these include the overall layout, boundaries of the environment, wall lengths, and angle dimensions (Cheng and Newcombe, 2005; Tommasi et al., 2012; Giocomo, 2016). Several studies in virtual environments have emphasized the idea that old age hinders the ability to use landmark information for navigation (Picucci et al., 2009; Harris et al., 2012; Wiener et al., 2012; Zhong and Moffat, 2016; Hartmeyer et al., 2017). Bécu and colleagues (2020) recently elaborated on such results and unveiled a specific age-related deficit for landmark-based compared with geometry-based navigation in ecological settings. Older participants were found to be impaired in anchoring their spatial behavior to landmark information and relied preferentially on geometric cues when both types of visual spatial cues were informative.

Despite the extensive body of literature characterizing the neural underpinnings of human spatial navigation (for recent reviews see Chersi and Burgess, 2015; Spiers and Barry, 2015; Epstein et al., 2017; Herweg and Kahana, 2018; Julian et al., 2018), few experiments have explored this question in the context of healthy aging. Indeed, only fifteen peer-reviewed neuroimaging studies have focused on spatial processing in normal aging and the majority of these studies have used structural analyses (Li and King, 2019). These studies highlight an age-related decline in place-based navigation associated with structural and functional changes to the hippocampus. A unique fMRI study has investigated the link between the use of visual spatial cues and the navigational skills of young and older adults (Schuck et al., 2015a). The authors combined computational modeling and fMRI during a virtual navigation task to examine how participants learned object locations relative to a circular enclosure or to a salient landmark. Young participants were found to use a hippocampal-dependent system for the representation of geometry (circular arena) and a striatal-dependent system for the representation of a landmark (traffic cone). It was further revealed that older participants relied on hippocampal structures for landmark-based navigation and that they were insensitive to geometric information provided by the environmental boundaries. This absence of reliance on geometric information is surprising considering the behavioral findings mentioned above, and could be related to the small field of view inherent to the scanner (Sturz et al., 2013).

Several other brains regions, known to be altered in healthy aging (Lester et al., 2017; Zhong and Moffat, 2018), have also been identified as key for the processing of relevant visual spatial cues for navigation (Epstein and Vass, 2014; Julian et al., 2018). Recently, there has been growing interest in unearthing the roles of the parahippocampal place area (PPA), the occipital place area (OPA), and the retrosplenial cortex (RSC), regions that respond to the presentation of visual scenes such as landscapes or urban environments. These scene-selective areas have been speculated to integrate incoming visual inputs with higher-level cognitive processes (for a review see Epstein et al., 2017; Julian et al., 2018). In brief, the PPA is sensitive to navigationally relevant cues (Janzen and van Turennout, 2004; Epstein, 2008) and may be implicated in the recognition of spatial context (Marchette et al., 2015); the OPA has been associated with the processing of local elements in scenes (Kamps et al., 2016) as well as with the representation of environmental boundaries (Julian et al., 2016); and the RSC is suggested to anchor heading information to local visual cues (for a recent review of RSC functions see Mitchell et al., 2018). Research exploring the neural activity within scene-selective regions in the context of aging is still in its infancy but evidence is accumulating for age-related alterations in these regions. Functional changes in the PPA have been linked to deficient processing of visual scenes (Ramanoël et al., 2015) and lower activations in the RSC of older adults have been associated with difficulties in switching between navigation strategies (Zhong and Moffat, 2018). The OPA’s progression as a function of age has not been fully characterized, but recent findings from the host laboratory have hinted at its preserved connectivity with other navigational brain structures in healthy aging (Ramanoël et al., 2019).

Although behavioral studies have provided some evidence for differences in the use of landmark cues across the lifespan, there is a clear paucity of research exploring the functional and neurobiological factors responsible for the deterioration of landmark information processing in older age. To address this caveat, the present study used fMRI to investigate to what extent healthy aging influences behavior and neural activity associated with landmark-based navigation. A second objective consisted in deciphering the role played by scene-selective regions (PPA, OPA, RSC) in age-related landmark-based navigation deficits.

## 2. Materials and methods

### 2.1 Participants

Among the 25 young adults and 21 older adults who completed the experiment, 4 older adults were excluded: 2 for a lack of task understanding and 2 for severe in-scanner motion (movements > 5 mm across trials). Overall, 25 young adults (18 males; 25.4 ± 2.7 years) and 17 older adults (7 males; 73.0 ± 3.9 years) were included in the analyses. The participants were part of the French cohort study *SilverSight* (~350 subjects) established in 2015 at the Vision Institute, Quinze-Vingts National Ophthalmology Hospital, Paris. The battery of clinical and functional examinations used to enroll participants comprised an ophthalmological and functional visual screening, a neuropsychological evaluation, an oculomotor screening, an audio-vestibular assessment as well as a static/dynamic balance examination. The neuropsychological evaluation included the Mini Mental State Examination (MMSE; Folstein et al., 1975) and computerized versions of the 3D mental rotation test (Vandenberg and Kuse, 1978), perspective-taking test (Kozhevnikov and Hegarty, 2001) and Corsi block-tapping task (Corsi, 1973). Older participants had a score of 24^1^ or higher on the MMSE. All subjects were right-handed, they had normal or corrected-to-normal vision, and they had no history of neurological or psychiatric disorders. Centration measurements and acuity were evaluated at least 2 weeks before the experimental session with a view to order MRI-compatible glasses for participants requiring visual correction (manufactured by *Essilor*). Participants gave their written informed consent to participate in the study, which was approved by the Ethical Committee “CPP Ile de France V” (ID_RCB 2015-A01094-45, CPP N°: 16122).

### 2.2 Virtual navigation task

#### 2.2.1 The virtual environment

The virtual navigation task was displayed on a MRI-compatible liquid crystal display monitor (NordicNeuroLab, Bergen, Norway) positioned at the head of the scanner bore. Participants viewed the screen (size: 69.84 cm (H) × 39.26 cm (V); pixels: 1920 × 1080) at a distance of 115 cm via a mirror fixed above the head-coil. The visible part of the screen subtended approximately 34 × 20 degrees of visual angle.

The virtual environment was programmed with Unity3D game engine (Unity Technologies SF; San Francisco, CA; https://unity.com/) and had participants navigate actively in a first-person perspective. The virtual environment was a three-arm maze (Y-maze) consisting of three corridors radiating out from a center delimited by homogenous wooden-like walls. Two configurations were designed. In the landmark condition all arms were 18 virtual meters (vm) long and equiangular. Three light gray-colored objects (a square, a triangle and a circle) were placed in front of each short wall at the center of the maze (Figure 1-A). In the control condition, the arms were 18 vm long and equiangular, and the maze was devoid of objects (Figure 1-B).

**Figure 1.**
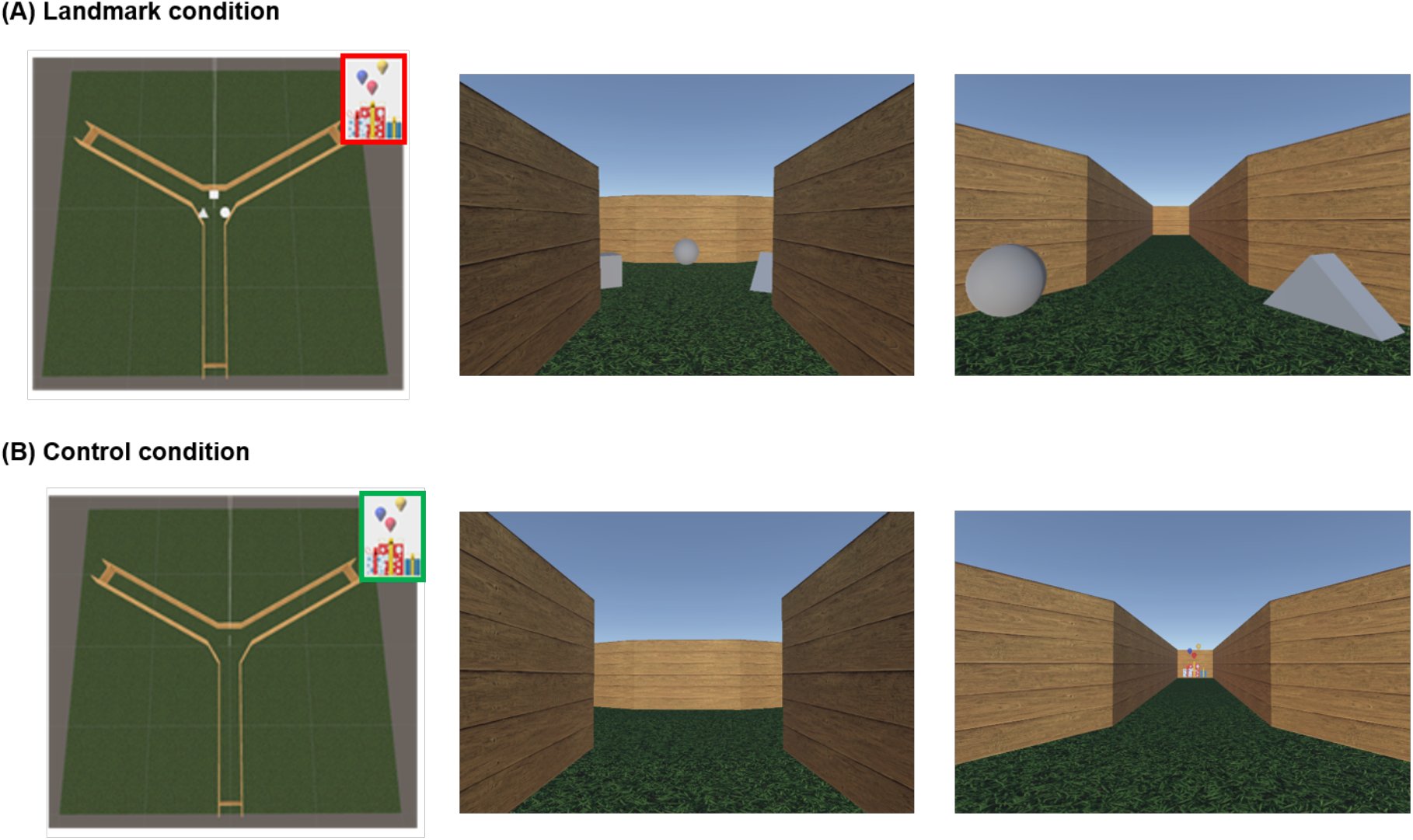
The virtual environment. **(A)** An overhead perspective of the environment for the landmark condition and two example views representing a first-person perspective within the maze. **(B)** An overhead perspective of the environment for the control condition and example views within the maze. Red: hidden goal; Green: visible goal. The aerial view was never seen by participants.

Participants navigated actively through the virtual environment with an MRI-compatible ergonomic two-grip response device (NordicNeuroLab, Bergen, Norway). They could move forward (thumb press), turn right (right index press) and turn left (left index press). A single finger press was necessary to initiate or stop movement. The forward speed of movement was set at 3 vm/s and the turning speed at 40°/s.

#### 2.2.2 Task design

Prior to scanning, all participants familiarized themselves with the response device in an unrelated virtual space both outside and inside the scanner. They were required to navigate within a square open-field environment and to walk over a wooden board that appeared at different locations.

The scanning session during the navigation task was divided into three runs: an encoding phase and a retrieval phase for the landmark condition and a control condition. At the start of the encoding phase participants were positioned in the center of the maze randomly facing one of the three arms. They were instructed to find a goal (gifts) hidden at the end of one corridor and remember its location using the visual information available in the center of the environment (the three light gray-colored objects). The encoding phase lasted 3 min to ensure that participants could explore all corridors. The retrieval phase in the same environment then began. In each trial participants were placed at the end of one of the two corridors that didn’t contain the goal with their back against the wall. The starting positions across trials were pseudo-randomized across subjects. Participants were asked to navigate to the previously encoded goal location. Upon arrival at the end of the correct arm, the gifts appeared to indicate successful completion of the trial and a fixation cross on a gray screen was presented for an inter-trial interval of 3-8 s. Participants needed to complete seven trials. The control condition consisted of a retrieval phase only; it was designed to account for potential confounding factors such as motor and simple perceptual aspects of the task. It was always performed last and it comprised four trials. Subjects started from the end of an arm and moved to the center of the maze from where the target was readily visible. They were instructed to navigate towards it. For both conditions, we recorded the trial duration and the response device use.

A short debriefing phase concluded the experimental session. Participants were probed on the strategy they used to orient in the landmark condition. They were asked to report how they solved the task: i) using one object, ii) using at least two objects, iii) randomly, iv) other strategy. Participants were deemed to be using a *place-based* strategy when their decision was based on two landmarks or more and to be using a *response-based* strategy when their decision was based on a single visual spatial cue (Iaria et al., 2003; Iglói et al., 2010, 2014; Chrastil, 2013; Gazova et al., 2013; Packard and Goodman, 2013; Colombo et al., 2017; Laczo et al., 2017). No participants answered that they oriented randomly or that they used a different strategy.

### 2.3 Functional localizer experiment

A block fMRI paradigm was used to locate scene-selective areas (Ramanoël et al., 2019). Scene-selective areas comprise the parahippocampal place area (PPA), the occipital place area (OPA) and the retrosplenial cortex (RSC). Participants were presented with blocks of 900 × 900 pixel grayscale photographs (18 × 18 degrees of visual angle) representing scenes, faces, everyday objects, and scrambled objects. The functional run lasted 4 min 40 s and it was composed of fourteen 20-s task blocks (4 blocks of scenes, 2 blocks of faces, 2 blocks of objects, 2 blocks of scrambled objects, and 4 blocks of fixation). Each stimulus was presented for 400 ms followed by a 600 ms inter-stimulus interval. Participants performed a “one-back” repetition detection task.

### 2.4 MRI Acquisition

Data were collected using a 3 Tesla Siemens MAGNETOM Skyra whole-body MRI system (Siemens Medical Solutions, Erlangen, Germany) equipped with a 64-channel head coil at the Quinze-Vingts National Ophthalmology Hospital in Paris, France. T2*-weighted echo-planar imaging (EPI) sequences, optimized to minimize signal dropout in the medial temporal region (Weiskopf et al., 2006), were acquired for functional imaging during the navigation task (voxel size = 3 × 3 × 2 mm, TR/TE/flip angle = 2685 ms/30 ms/90°, interslice gap = 1 mm, slices = 48, matrix size = 74 × 74, FOV = 220 × 220 mm). For the localizer experiment, 284 volumes from 64 slices were acquired using a T2*-weighted simultaneous multi-slice echo planar sequence (SMS-EPI; voxel size = 2.5 × 2.5 × 2.4 mm, TR/TE/flip angle = 1000 ms/30 ms/90°, matrix size = 100 × 100, SMS = 2, GRAPPA = 2). The anatomical volume consisted of a T1-weighted, high-resolution three-dimensional MPRAGE sequence (voxel size = 1 × 1 × 1.2 mm, TR/TE/IT/flip angle = 2300 ms/2.9 ms/900 ms/9°, matrix size = 256 × 240 × 176).

### 2.5 Statistical analyses of behavioral data

Normality of data was assessed graphically with quantile-quantile plots and numerically with the Shapiro-Wilk test. Descriptive characteristics and cognitive measures were compared across age groups using independent samples t-test for normally distributed continuous data, Mann-Whitney U tests for non-normally distributed continuous data and chi-square test of independence for categorical data. Navigation performance (time to reach the goal, error rate) was compared across age group and sex with Mann-Whitney U tests. A logistic regression adjusted for sex was conducted to examine the relationship between age and strategy use in the landmark condition.

### 2.6 Preprocessing and statistical analyses of fMRI data

#### 2.6.1 Whole brain analyses

FMRI data analysis was performed using a combination of SPM12 release 7487 (Wellcome Department of Imaging Neuroscience, London, UK) and ArtRepair toolbox (Mazaika et al., 2009) implemented in MATLAB 2015 (Mathworks Inc., Natick, MA, USA). The first five functional volumes of the encoding, retrieval and control runs were discarded to allow for equilibration effects. Slice-timing correction was applied and functional images were realigned to the mean functional image using a rigid body transformation. Artefacts related to motion were then examined with ArtRepair. Two older subjects were subsequently excluded. Volumes displaying elevated global intensity fluctuation (> 1.3%) and movement exceeding 0.5 mm/TR were repaired using interpolation from adjacent scans. The T1-weighted anatomical volume was then realigned to match the mean functional image of each participant and normalized to the Montreal Neurological Institute (MNI) space using a 4^th^ degree B-Spline interpolation. The anatomical normalization parameters were subsequently used for the normalization of functional volumes. Each functional scan was smoothed with an 8 mm FWHM (Full Width at Half Maximum) Gaussian kernel. The preprocessed images were visually inspected to ensure that there were no realignment or normalization issues.

Statistical analysis was performed using the general linear model for block design at the single participant level (Friston et al., 1995). The seven trials of the retrieval phase in the landmark condition, the four trials of the control condition and fixation times were modeled as regressors, constructed as box-car functions and convolved with the SPM hemodynamic response function (HRF). The encoding phase was not included in the analysis as its duration differed greatly between participants. Time to reach the goal by trial, grip response during navigation, and movement parameters derived from the realignment correction (three translations and three rotations) were entered in the design matrix as regressors of no-interest. Time series for each voxel were high-pass-filtered (1/128 Hz cutoff) to remove low-frequency noise and signal drift. Individual contrasts were submitted to a multiple regression and a two-samples t-test. Sex and total brain volume were included as covariates in the regression and total brain volume was included as a covariate in the two-samples t-test (see section 3.1). Areas of activation were tested for significance using a statistical threshold of *p* < 0.001 uncorrected at voxel-level, with a minimum cluster extent of k = 10 voxels (Iglói et al., 2010, 2014; Sutton et al., 2010; Schuck et al., 2015a; Javadi et al., 2017).

#### 2.6.2 Region of interest analyses

Data from the localizer experiment were analyzed using SPM12. For each participant, the first 4 functional localizer volumes were discarded and the remaining images were realigned, co-registered to the T1-weighted anatomical image, normalized to the MNI space and smoothed using an 8 mm FWHM Gaussian kernel. Slice-timing correction was not applied following recommendations from the Human Connectome Project functional preprocessing pipeline for multi-slice sequences (Glasser et al., 2013). The localizer images were analyzed using a single participant general linear model for block design. Five conditions of interest (scenes, faces, objects, scrambled objects, fixation) were modeled as five regressors and convolved with a canonical HRF. Movement parameters were included in the model as regressors of no interest and each voxel’s time-series was high-pass-filtered (1/128 Hz cutoff).

PPA, OPA and RSC regions were located independently for each participant using the fMRI contrast [Scenes > (Faces + Objects)]. Significant voxel clusters on individual t-maps were identified using family-wise error correction (FWE) for multiple comparisons (alpha = 0.05). Mask ROIs were created as the 40 contiguous voxels with the highest t-values around the peaks of activation from the left and right hemispheres. The two 40-voxel regions from each hemisphere were subsequently summed into a single 80-voxel ROI. Mean parameter estimates for the contrast [Landmark > Control] were extracted from the three mapped ROIs using the REX MATLAB-based toolkit. The mean values from each ROI were compared between young and older subjects using two samples t-test and significance was set at *p* < 0.0083 after Bonferroni correction for multiple comparisons (p = 0.05/(3 × 2)).

## 3. Results

### 3.1 Behavioral results

Data from 25 young and 17 older participants were analyzed. Neuropsychological assessments showed that older adults had significantly poorer performance than young adults across all measures. Descriptive and cognitive characteristics are summarized in Table 1.

**Table 1.** Descriptive characteristics and cognitive performance of young and older participants. M: male; F: female; SEM: standard error of the mean; MMSE: mini mental state examination; ^1^Independent samples t-test; ^2^Mann Whitney U test.

| Sex (M/F) | Groups |  | p-value |
| --- | --- | --- | --- |
|  | Young adults | Older adults |  |
|  | 18/7 | 7/10 |  |
| | Mean (± SEM) | Mean (± SEM) | |
| Age<sup>1</sup> | 25.4 (± 0.5) | 73.0 (± 0.9) | p < 0.001 |
| Total brain volume<sup>1</sup> (cm<sup>3</sup>) | 1301 (± 18) | 1061 (± 23) | p < 0.001 |
| MMSE<sup>2</sup> | 30.0 (± 0.0) | 28.8 (± 0.4) | p < 0.001 |
| 3D mental rotation<sup>1</sup> | 18.3 (± 0.9) | 12.7 (± 1.2) | p < 0.001 |
| Corsi forward<sup>2</sup> | 7.2 (± 0.2) | 4.4 (± 0.2) | p < 0.001 |
| Corsi backward<sup>2</sup> | 6.2 (± 0.3) | 4.6 (± 0.2) | p < 0.001 |
| Perspective taking test<sup>2</sup> | 15.3 (± 1.7) | 46.1 (± 6.7) | p < 0.001 |

Navigation performance across age groups is presented in Figure 2. We found that older adults chose the wrong corridor in 10% of trials while young adults made no errors (X² (1, N = 42) = 12.35, *p* = 0.006; Figure 2A). In addition, older subjects were significantly slower to reach the goal in the landmark condition than younger subjects (mean ± SEM: 19.85 s ± 1.67 vs 11.97 s ± 0.13; U(40) = 5.267, *p* < 0.001; Figure 2B). There was no sex effect on navigation time, defined by the average time to reach the goal, in each age group separately. However, when data from both age groups were pooled women’s navigation time appeared to be significantly longer than men’s (17.75 s ± 1.88 s vs 13.40 s ± 0.64 s; U(40) = 2.294, *p* = 0.022). Sex and total intracranial volume were therefore included as covariates in the fMRI multiple regression analyses. We further found that age was a significant predictor of strategy use (R^2^ = 0.967, *p* = 0.048). Older adults were less likely to rely on a place-based strategy during landmark-based navigation than younger adults (Figure 2C).

**Figure 2.**
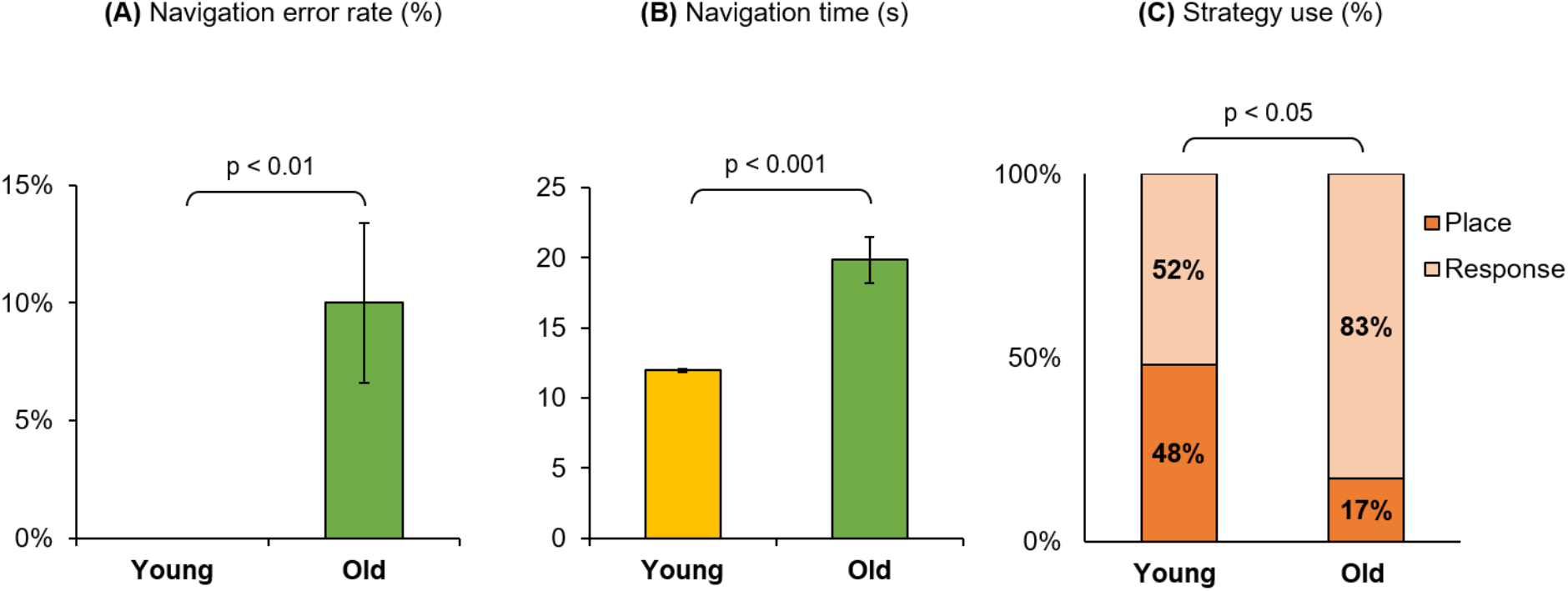
Behavioral results for the virtual navigation task across age groups. **(A)** Proportion of trials in which the wrong corridor was chosen (navigation error rate). **(B)** Time taken to reach the goal averaged across 7 trials in the landmark condition (navigation time). **(C)** Proportion of participants who used a place-based or response-based strategy in the landmark condition (strategy use). Error bars represent the standard errors of the mean.

### 3.2 Whole-brain

#### 3.2.1 Multiple regression analyses

To locate the brain regions related to navigation performance, we first examined the association between both groups’ navigation time and patterns of brain activity for the fMRI contrast [Landmark > Control]. We observed a negative association between navigation time and neural activity in many clusters across the brain (Table 2 and Figure 3) including frontal (right superior and middle gyri), temporal (middle, inferior, lingual and parahippocampal gyri), parietal (left angular gyrus including the inferior parietal lobule) and occipital cortices (left superior occipital gyrus) as well as in the cerebellum (lobule VI and vermis). Temporal activations in the left hemisphere comprised the posterior part of the hippocampus: CA1 and the Dentate Gyrus (x=−24, y=−46, z=5 and x=−36, y=−43, z=−4). Other activations included the ventral temporal cortex (x=45, y=8, z=−37 and x=54, y=5, z=−34) and the visual area V3A (x=−21, y=−97, z=23) that is part of the dorsal visual stream. The inverse association did not elicit any significant brain activations. Complementary analyses revealed a negative association between age and patterns of neural activity in the left superior temporal gyrus only (x=−45, y=−1, z=−16) for the fMRI contrast [Landmark > Control]. The latter finding suggests that the negative association between navigation time and temporal activations previously reported is largely driven by age.

**Table 2.**
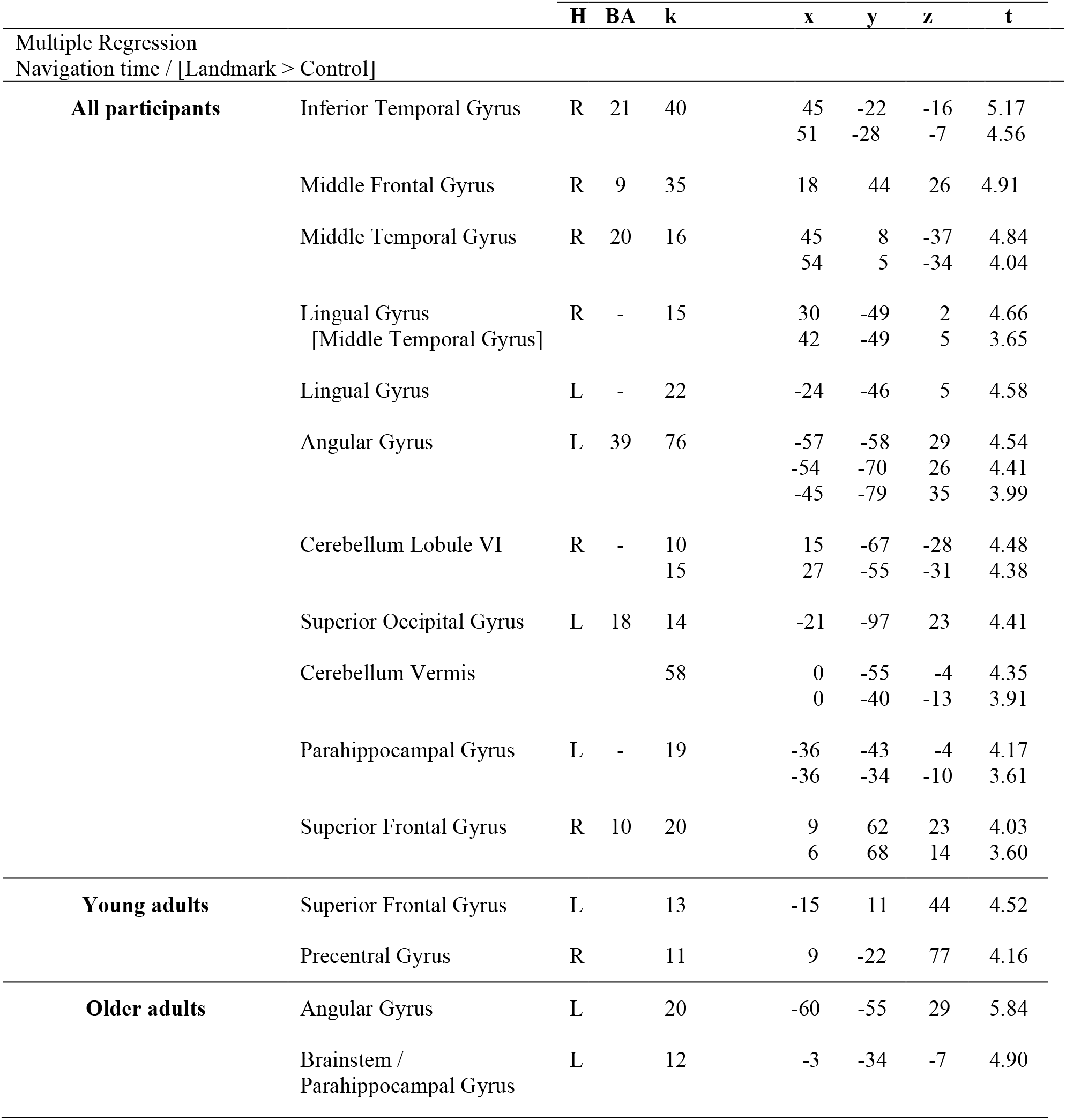
Cerebral regions whose activity for the contrast [Landmark > Control] was predicted by navigation time across all participants and across age groups (sex and intracranial volume were included as covariates). The statistical threshold was defined as *p* < 0.001 uncorrected for multiple comparisons at voxel-level with an extent voxel threshold set at 10 voxels. For each cluster, the region with the maximum t-value is listed first and other regions in the cluster are listed underneath [in square brackets]. Montreal Neurological Institute (MNI) coordinates (x, y, z) of the peak activation and number of voxels (k) in a cluster are also shown. H = hemisphere; R = right hemisphere; L = left hemisphere; BA = Brodmann area.

**Figure 3.**
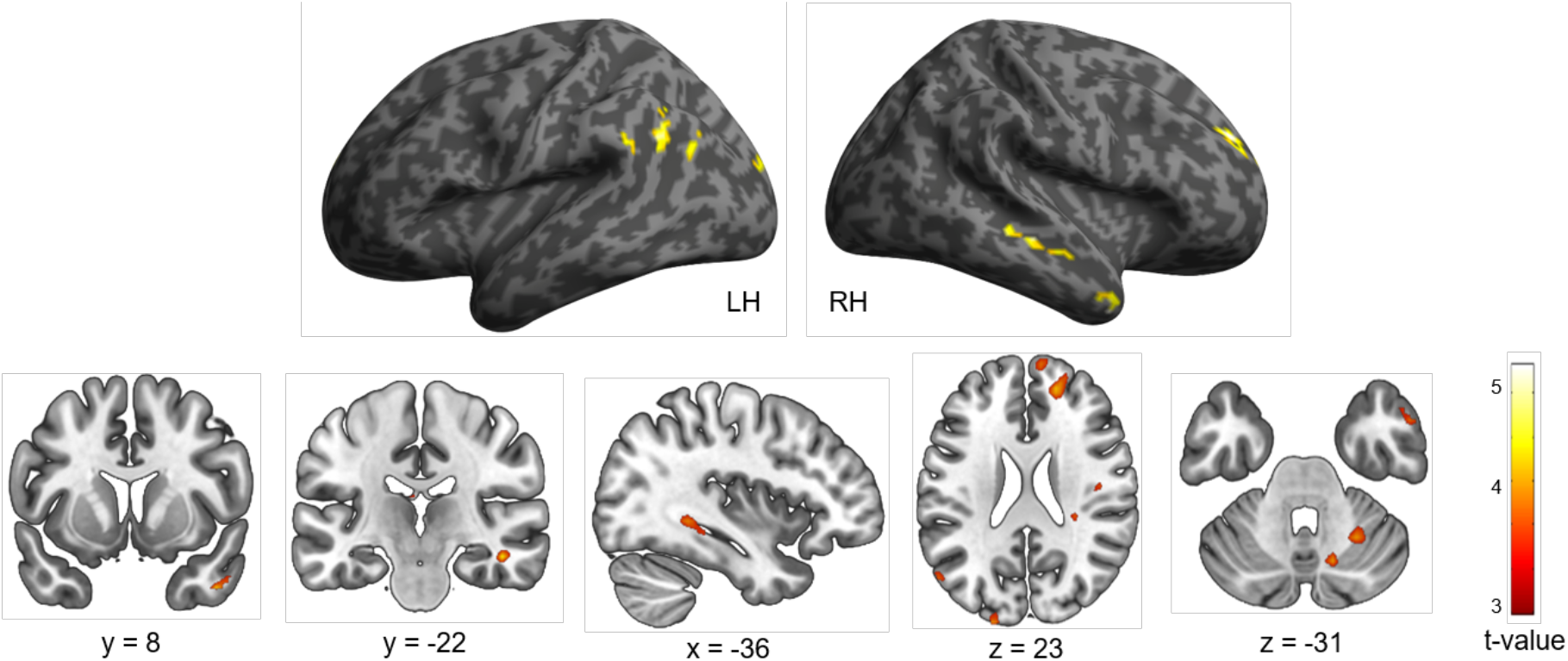
Cerebral regions whose activity for the contrast [Landmark > Control] was predicted by navigation time projected onto 3D inflated anatomical templates and 2D slices (*p* < 0.001 uncorrected, k = 10 voxels). LH: left hemisphere; RH: right hemisphere.

We then investigated the relationship between navigation time and neural activity in each age group separately (Table 2). In the young participant group, navigation performance was associated with neural activity in the left superior frontal gyrus and in the right precentral gyrus. In the older participant group, results revealed a main cluster in the left angular gyrus, including the inferior parietal lobule, as well as a small cluster encompassing the brainstem and the parahippocampal gyrus.

#### 3.2.2 Two-sample analyses

Results for the within-group and between-group analyses are shown in Table 3 and Figure 4. We first contrasted brain activity during the landmark condition with activity during the control condition [Landmark > Control] for each age group individually. For young participants, we found significant clusters in the superior and inferior temporal gyri of the left hemisphere, which included the amygdala and hippocampus, in the middle and inferior occipital gyri bilaterally and in the right cerebellum (Crus I-II). In the older participant group, the fMRI contrast [Landmark > Control] did not elicit any significant activation. Direct comparisons between age groups revealed significant results for the fMRI contrast [Young > Old] but not for the contrast [Old > Young]. Young participants exhibited stronger activity in the left inferior temporal gyrus, comprising the amygdala and hippocampal regions, than older subjects.

**Table 3.**
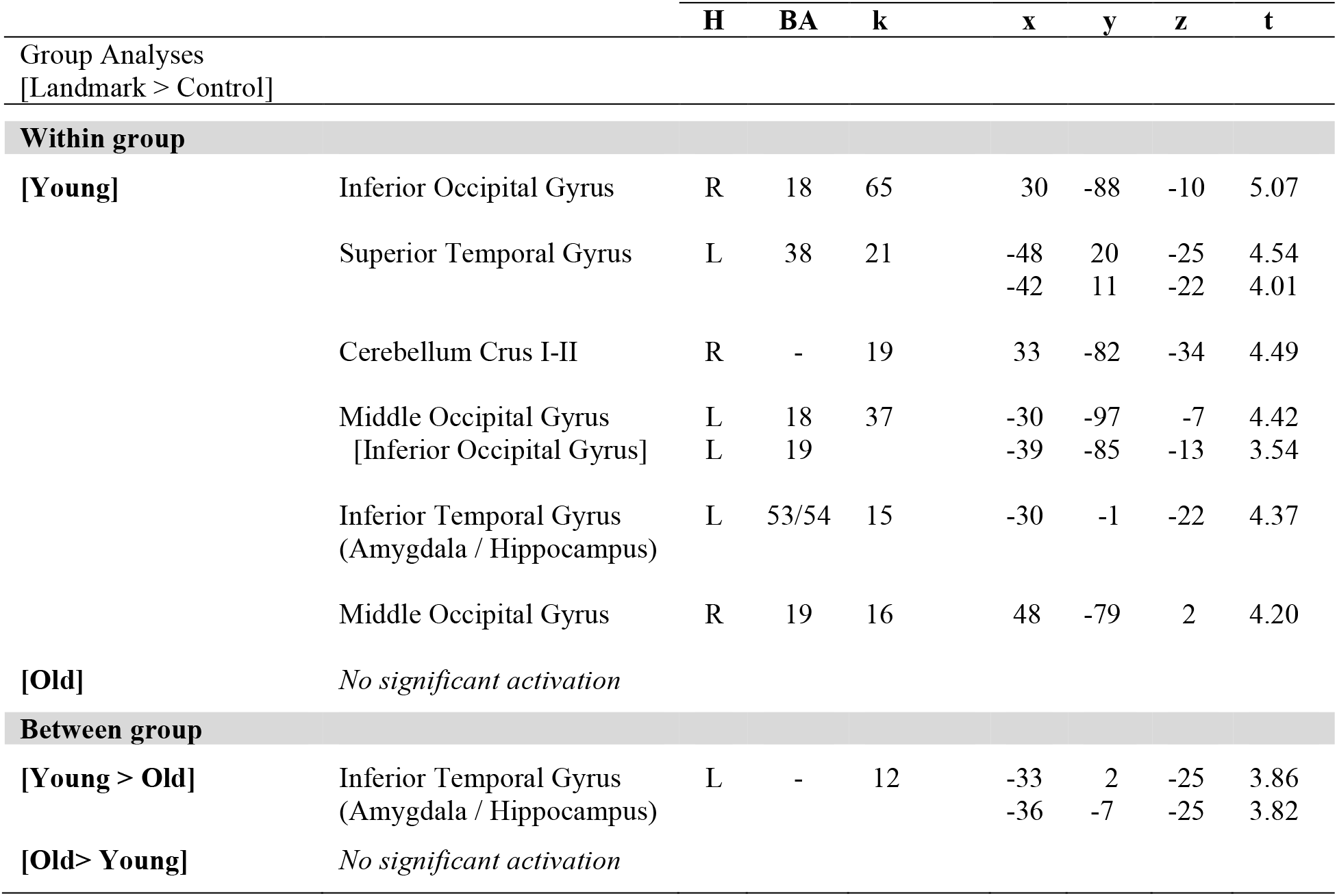
Cerebral regions whose activity for the contrast [Landmark > Control] was elicited by within-group or between-group analyses (total intracranial volume was included as a covariate). The statistical threshold was defined as *p* < 0.001 uncorrected for multiple comparisons at voxel-level with an extent voxel threshold set at 10 voxels. For each cluster, the region with the maximum t-value is listed first and other regions in the cluster are listed underneath [in square brackets]. Montreal Neurological Institute (MNI) coordinates (x, y, z) of the peak activation and number of voxels (k) in a cluster are also shown. H = hemisphere; R = right hemisphere; L = left hemisphere; BA = Brodmann area.

**Figure 4.**
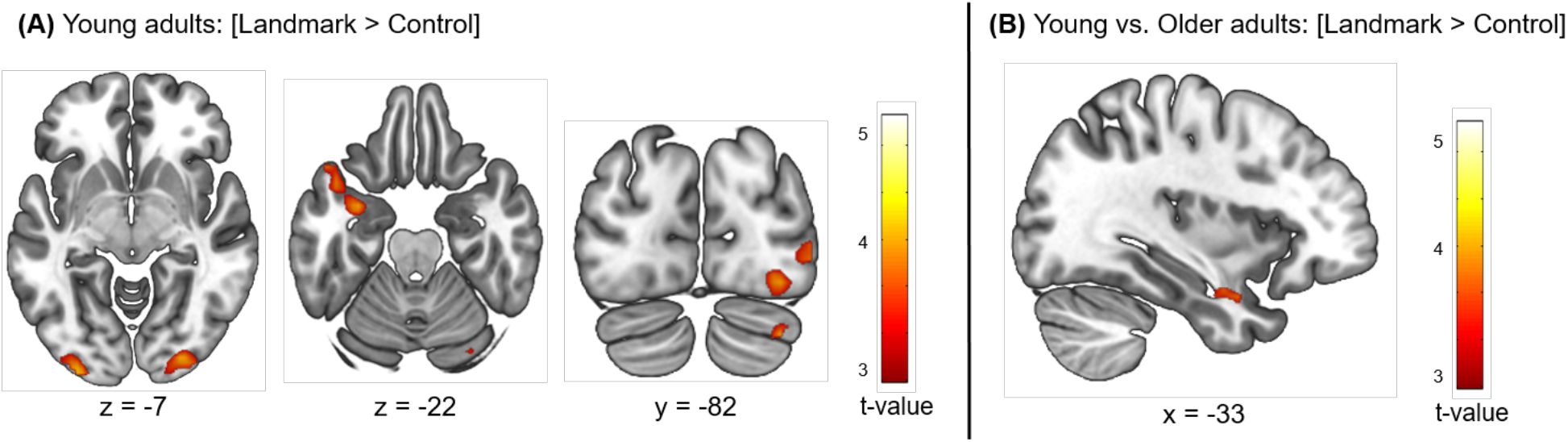
Cerebral regions whose activity for the contrast [Landmark > Control] was elicited by within-**(A)** or between-group **(B)** analyses projected onto 2D slices (*p* < 0.001 uncorrected, k = 10 voxels).

The absence of activation in the older adult group prompted us to conduct further analyses. We first investigated the cognitive cost of the control condition in each age group by comparing cerebral activations for the fMRI contrast [Control > Fixation]. The group comparison [Young > Old] showed no significant activation. However, the inverse group comparison [Old > Young] revealed extended activations in the right precuneus, the right superior temporal gyrus, the right supramarginal gyrus and superior parietal lobule as well as several bilateral frontal regions (Table 4). These results hint at the possibility that the control condition was cognitively more demanding for older participants than for young participants. We performed a second group comparison with the fMRI contrast [Landmark > Fixation] using a conservative cluster-level FWE correction *p* < 0.05. Significant activations were reported for the [Old > Young] comparison only, and included the middle frontal and superior parietal gyri in each hemisphere, the left supramarginal gyrus as well as the right cerebellum (Table 5).

**Table 4.** Cerebral regions whose activity for the contrast [Control > Fixation] was elicited by between-group analyses (total intracranial volume was included as covariate). The statistical threshold was defined as p < 0.001 uncorrected for multiple comparisons at voxel-level with an extent voxel threshold set at 10 voxels. For each cluster, the region with the maximum t-value is listed first and other regions in the cluster are listed underneath [in square brackets]. Montreal Neurological Institute (MNI) coordinates (x, y, z) of the peak and number of voxels (k) of clusters are also shown. CTRL = Control condition; Fix = Fixation; H = hemisphere; R= right hemisphere; L = left hemisphere; BA = Brodmann area.

|  |  | H | BA | k | x | y | z | t |
| --- | --- | --- | --- | --- | --- | --- | --- | --- |
| Group Analyses<br>[CTRL > Fix] |  |  |  |  |  |  |  |  |
| [Young > Old] | No significant activation |  |  |  |  |  |  |  |
| [Old > Young] | Middle Frontal Gyrus | L | 8 | 80 | -27 | 32 | 56 | 5.56 |
|  | [Middle Frontal Gyrus] |  |  |  | -36 | 23 | 56 | 4.58 |
|  | [Superior Frontal Gyrus] |  | 6 |  | -18 | 32 | 62 | 4.05 |
|  | Superior Temporal Gyrus | R | - | 18 | 36 | 17 | -19 | 4.71 |
|  | Superior Frontal Gyrus | R | 10 | 32 | 12 | 56 | -10 | 4.56 |
|  | [Middle Frontal Gyrus] |  |  |  | 15 | 44 | -4 | 4.02 |
|  | Inferior Temporal Gyrus | L | 37 | 25 | -54 | -49 | -13 | 4.39 |
|  | Supramarginal Gyrus | R | 40 | 31 | 42 | -40 | 41 | 4.23 |
|  | [Superior Parietal Gyrus] |  | 7 |  | 33 | -46 | 38 | 4.21 |
|  | Superior Frontal Gyrus | L | 32 | 14 | -12 | 41 | 2 | 4.16 |
|  | [Middle Frontal Gyrus] |  |  |  | -18 | 41 | -4 | 4.10 |
|  | Inferior Frontal Gyrus | L | 46 | 38 | -36 | 35 | 14 | 4.15 |
|  |  |  |  |  | -48 | 41 | 11 | 3.85 |
|  | Precuneus | R | 31 | 12 | 9 | -58 | 38 | 4.00 |
|  | Middle Frontal Gyrus | L | 8 | 20 | -51 | 20 | 41 | 3.92 |
|  |  |  |  |  | -48 | 11 | 50 | 3.81 |
|  | Inferior Frontal Gyrus | L | 45 | 12 | -57 | 23 | 8 | 3.79 |

**Table 5.** Cerebral regions whose activity for the contrast [Landmark > Fixation] was elicited by between-group analyses (total intracranial volume was included as covariate). The statistical threshold for cluster was defined as p < 0.05 FWE-corrected for multiple comparisons with an extent voxel threshold set at 10 voxels. For each cluster, the region with the maximum t-value is listed first and other regions in the cluster are listed underneath [in square brackets]. Montreal Neurological Institute (MNI) coordinates (x, y, z) of the peak and number of voxels (k) of clusters are also shown. LMK = Landmark condition; Fix: Fixation condition; H = hemisphere; R = right hemisphere; L = left hemisphere; BA = Brodmann area; FWE = family-wise error.

|  |  | H | BA | k | x | y | z | t |
| --- | --- | --- | --- | --- | --- | --- | --- | --- |
| Group Analyses<br>[LMK > Fix] |  |  |  |  |  |  |  |  |
| [Young > Old] | No significant activation |  |  |  |  |  |  |  |
| [Old > Young] | Middle Frontal Gyrus | R |  | 22 | 24 | 47 | -1 | 6.06 |
|  | Angular Gyrus | L |  | 223 | -30 | -64 | 47 | 5.66 |
|  | [Superior Parietal Gyrus] |  |  |  | -33 | -55 | 44 | 5.59 |
|  | [Supramarginal Gyrus] |  |  |  | -48 | -43 | 44 | 5.32 |
|  | Middle Frontal Gyrus | R |  | 34 | 39 | 38 | 17 | 5.37 |
|  |  |  |  |  | 33 | 47 | 20 | 4.47 |
|  | Cerebellum | R | - | 23 | 36 | -73 | -22 | 5.19 |
|  | Middle Frontal Gyrus | L |  | 127 | -42 | 5 | 41 | 5.14 |
|  |  |  |  |  | -45 | 26 | 26 | 5.13 |
|  |  |  |  |  | -45 | 5 | 50 | 5.12 |
|  | Superior Parietal Gyrus | R |  | 36 | 30 | -55 | 44 | 4.89 |
|  | [Angular Gyrus] |  |  |  | 30 | -64 | 53 | 4.64 |

### 3.3 Regions of interest

The PPA, OPA and RSC ROIs were defined for each individual based on the independent localizer experiment. We first examined age-related differences in average parameter activity for the fMRI contrast [Landmark > Control]. No significant differences were found. We conducted a second analysis based on the fMRI contrast [Landmark > Fixation]. The group comparison revealed significantly enhanced OPA activity in older adults compared with young adults after adjusting for multiple comparisons (mean ± SEM: 2.19 ± 0.27 vs. 1.07 ± 0.21; t = −3.4, *p* = 0.002). No significant differences were observed in the activity of the PPA and RSC between young and older participants (Figure 5).

**Figure 5.**
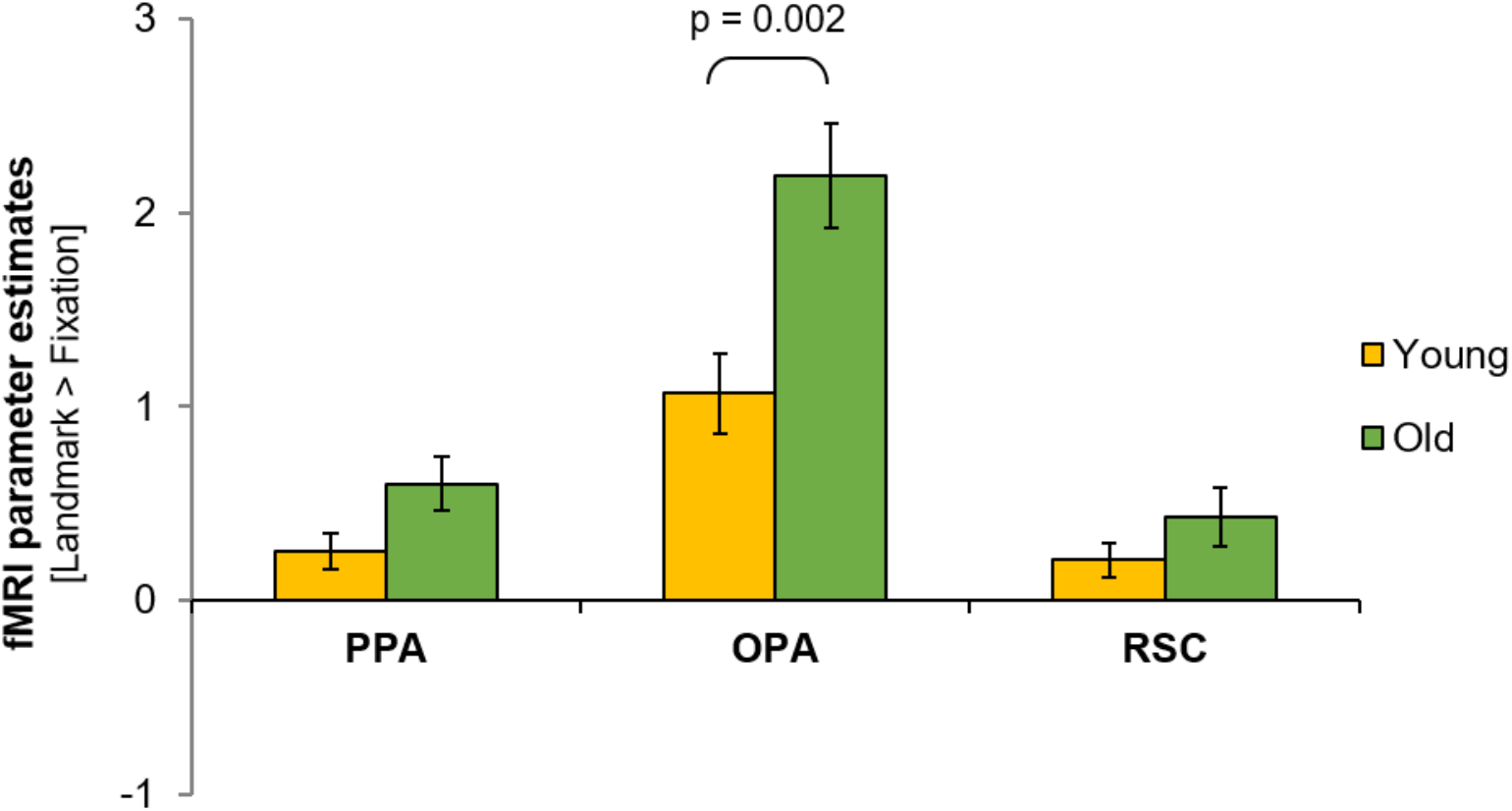
FMRI parameter estimates in the PPA, OPA, and RSC for the fMRI contrast [Landmark > Fixation] across age groups. Error bars reflect standard errors of the mean.

## 4. Discussion

The present fMRI study examined age-related differences in landmark-based navigation using a non-ambiguous Y-maze reorientation paradigm. The task was designed to limit the influence of mnemonic and motor components in order to gain a specific understanding of the neural bases subtending visual spatial cue reliance in young and healthy older adults.

### Behavior

We first replicated well-established findings and showed that older adults had lower scores than their younger counterparts on visuo-spatial cognitive neuropsychological measures, including the perspective-taking, 3D mental rotation and Corsi block-tapping tasks (Ohta et al., 1981; Clancy Dollinger, 1995; Iachini et al., 2005; Techentin et al., 2014). These tests are known to be good predictors of general navigation skills and their decline with age may account in part for older adults’ deficient navigation performance (Zhong and Moffat, 2016). Notably, perspective-taking, mental rotation, and spatial memory are important abilities for spatial learning and for the dynamic manipulation of sensory information during navigation (Allen et al., 1996; Kozhevnikov et al., 2006; Meneghetti et al., 2018; Muffato et al., 2020).

Consistent with past literature, we found that older subjects’ navigation performance was significantly poorer than young subjects’ and we reported a bias for response-based strategies in older adults (Moffat and Resnick, 2002; Harris and Wolbers, 2012; Rodgers et al., 2012; Gazova et al., 2013; Wiener et al., 2013; Schuck et al., 2015b; van der Ham et al., 2015; Zhong and Moffat, 2016; Kimura et al., 2019; Merhav and Wolbers, 2019; Bécu et al., 2020). This navigation strategy preference appeared to be associated with older age. However, we cannot exclude the possibility that the differential proportion of women in the two age groups (young: 28% vs. old: 59%) partly accounted for the increased use of response-based strategies in older adults (Perrochon et al., 2018). Notwithstanding these age-related differences, it is important to mention that older participants achieved a high level of performance on the task and made few errors. We argue that this result stemmed from the simplicity of the virtual environment that contained a unique junction and three proximal landmarks (Moffat and Resnick, 2002; Caffo et al., 2018). Moreover, both place- and response-based strategies could be used to successfully complete the task.

### Whole-brain

In accordance with previous neuroimaging studies looking at the neural bases of spatial navigation, we found that landmark-based navigation recruited an extended network of brain regions (Kuhn and Gallinat, 2014; Spiers and Barry, 2015b; Coughlan et al., 2018; Cona and Scarpazza, 2019).

This network spanned posterior structures linked to visuo-spatial processing. We reported activation of the left superior occipital gyrus which corresponds to visual area V3A and is involved in optic flow tracking for visual path integration (Sherrill et al., 2015; Zajac et al., 2019). In addition, our landmark-based navigation paradigm elicited activity in the ventral temporal cortex. The latter is known to process high-level visual information such as object quality (Kravitz et al., 2013; Nau et al., 2018). The recruited network also encompassed the posterior section of the hippocampus and the parahippocampal gyrus; brain areas that play a central role in spatial navigation and that are particularly active during immediate retrieval phases of navigation paradigms (Kuhn and Gallinat, 2014; Cona and Scarpazza, 2019). Furthermore, we found significant activity in the angular gyrus, a region of the posterior parietal cortex known to encode landmarks in the environment with respect to the self (Ciaramelli et al., 2010; Auger and Maguire, 2018). Our task also prompted activation of the prefrontal cortex which is thought to contribute to spatial working memory during active navigation (Wolbers and Hegarty, 2010; Ito, 2018). It thus appears that accurate landmark-based navigation required the integration of objects within a first-person framework and the maintenance of such representations in working memory (Sack, 2009; Seghier, 2012; Miniaci and De Leonibus, 2018). Finally, we found that lobule VI and the vermis of the right cerebellum were active during landmark-based navigation. This finding is in accordance with the cerebellum’s postulated role in cognitive aspects of spatial navigation (Rochefort et al., 2013). We must nonetheless acknowledge the eventuality that cerebellar activity reflected sensory-motor processing such as the degree of motor learning or eye and finger movements (Bo et al., 2011; Igloi et al., 2015).

Importantly, older participants were found to over-recruit superior parietal regions compared to young participants. One could speculate that the increased bilateral activity in the angular gyrus drove the encoding of landmarks in a first-person perspective leading to a bias for response strategies in the older adult group. This is consistent with the observed age-related reduction in temporal activity. Indeed, changes in strategy preference with advancing age have been extensively documented and they are thought to be mediated by a shift from the hippocampal regions towards other cerebral structures such as the parietal cortex (Rodgers et al., 2012; Wiener et al., 2013). Within this framework of interpretation, older adults’ increased cerebellar activity could also reflect a change in strategy preference as recent evidence has implicated the cerebellum in the mediation of response-based strategies (Igloi et al., 2015). Older participants further displayed enhanced activation of frontal cortices. Various authors have stressed the impact of age-related modifications in the prefrontal cortex on hippocampal and striatal dynamics, which could contribute to impaired strategy implementation and switching (Lester et al., 2017; Goodroe et al., 2018; Zhong and Moffat, 2018). In contrast to previous studies that reported striatal activity during response-based navigation, our results did not show increased striatal activity in the older adult group (Konishi et al., 2013; Schuck et al., 2015b). Such a difference may be explained by the high proportion of young adults using response-based strategies in our task. Worthy of note, the differential patterns of neural activity observed in the young and older participant groups may be partially due to age-related cognitive and motor differences. Although we tailored the duration of the familiarization phase to each subject’s needs and controlled for response device use, we cannot omit the potential influence of older adults’ lesser familiarity with new technologies and declining executive functions. For example, the lack of activity elicited by the contrast [Landmark > Control] in older subjects could reflect the deficient integration of new instructions when switching between tasks (Hirsch et al., 2016).

Of interest, whole-brain analyses revealed that young adults recruited the cortical projections of the central visual field in posterior occipital regions (Figure 6, MNI coordinates left: x=−24, y=−49, z=2 and right: x=30, y=−49, z=2). The latter brain area is dedicated to fine-grained visual perception such as object recognition (Wandell et al., 2005; Kauffmann et al., 2014). Exploratory between-group analyses showed that the fMRI contrast [Landmark] elicited more activity in anterior occipital regions (MNI coordinates x=6, y=−73, z=−1) that respond to the peripheral visual field in older participants than in young participants (Figure 6). Previous neuroimaging studies have uncovered specific age-related changes in the occipital regions associated with central visual field processing and a relative preservation of areas associated with peripheral visual field processing (Brewer and Barton, 2012; Ramanoël et al., 2015). Additionally, group comparisons revealed that young subjects had more activity in the anterior section of the inferior temporal gyrus than older subjects. As mentioned previously, the anterior temporal cortex is critical for perceptual recognition and visual object processing (Litman et al., 2009). Our findings are therefore consistent with recent evidence highlighting deficient fine-grained processing of sensory information in older adults and emphasize the importance of acute object discrimination for landmark-based navigation and episodic memory (Burke et al., 2018; Greene and Naveh-Benjamin, 2020). Taken together, the above results suggest that occipital and temporal regions involved in the representation of fine-grained information are particularly disrupted in older age. Further research is warranted to determine whether the age-related decline in orientation skills could stem from the less efficient processing of visual spatial cues.

**Figure 6.**
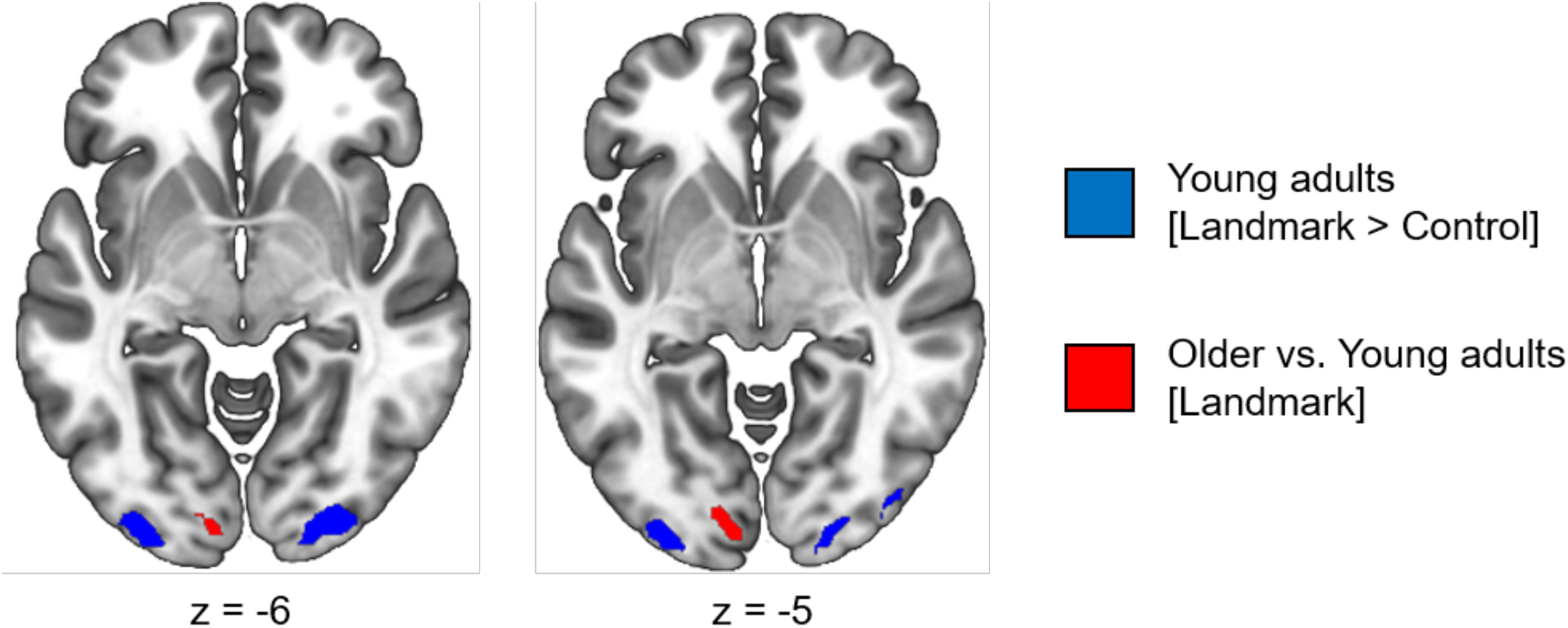
Occipital clusters recruited during the navigation task. In blue, the cortical projection of the central visual field that was elicited by the fMRI contrast [Landmark > Control] in young adults. In red, the cortical projection of the peripheral visual field that was elicited by comparing young and older adults for the fMRI contrast [Landmark].

### Scene-selective regions

Given the predominant role of visual perception in human spatial navigation (Ekstrom, 2015; Nau et al., 2018), there has been heightened interest in the PPA, RSC and OPA and their respective contributions to landmark processing (Epstein et al., 2017; Julian et al., 2018). Our results pointed to greater OPA activity in older participants compared with younger participants which is in line with recent work showing a higher functional connectivity around the OPA in older adults (Ramanoël et al., 2019).The OPA is known to be sensitive to self-perceived distance and motion (Persichetti and Dilks, 2016) and to extract the navigational affordances of the local visual scene from a first-person perspective (Bonner and Epstein, 2017). Critically, these self-centered navigation skills are relatively well preserved in healthy aging (Moffat, 2009). In line with the over-activation of the parietal cortex in older adults, one could conceive that the increased OPA activation in the older adult group reflects a compensatory mechanism to offset the reduced activity in the temporal cortex, thus mitigating age-related place learning deficits. As a side note, considering that the OPA has been causally linked to the processing of environmental boundaries (Julian et al., 2016), our result offers a potential explanation for older adults’ preferential reliance on geometric information in an ecological cue conflict paradigm (Bécu et al., 2020).

Surprisingly, we did not find differences in the activity of the RSC across age groups. Previous work has demonstrated an age-related decline in RSC activation during spatial navigation tasks (Meulenbroek et al., 2004; Moffat et al., 2006; Antonova et al., 2009). The RSC is known to mediate several cognitive functions pertaining to spatial navigation (Vann et al., 2009; Mitchell et al., 2018) including translation between reference frames and recollection of visual landmarks (Auger et al., 2015). The discrepancy between our results and those from the literature could be explained by the relative simplicity of our task. In contrast to previous research conducted with young adults, our paradigm strived to restrict mnemonic processing and comprised only three stable, salient and simple landmarks located at a single intersection (Wolbers and Büchel, 2005; Auger et al., 2012; Auger and Maguire, 2018). Another probable explanation lies in the idea that functional and structural changes to the RSC have been proposed to be more pronounced in pathological aging than normal aging (Fjell et al., 2014; Dillen et al., 2016).

Finally, we reported a weak recruitment of the PPA during landmark-based navigation with no significant difference across age groups. Previous studies have found the PPA to be involved in encoding the navigational relevance of objects for orientation (Janzen and van Turennout, 2004) and in landmark recognition (Epstein and Vass, 2014). As previously noted, our virtual environment comprised a small number of simple and non-ambiguous objects, the lack of activity in the PPA is thus unsurprising. In addition, two recent studies de-emphasized PPA’s contribution to active navigation and highlighted its specificity for place recognition (Persichetti and Dilks, 2018, 2019). Research exploring the neural activity within scene-selective regions in the context of aging is still in its infancy. Future studies are needed to better characterize age-related changes in brain areas implicated in processing both the visual and cognitive properties of spatial cues.

### Limitations

The current study has several limitations. First, we focused exclusively on the retrieval phase as the encoding phase proved to be too heterogeneous across participants. It would be of immediate interest to assess the influence of various visuo-perceptual modulations, such as the visibility of spatial cues, on the quality of spatial encoding. Second, although spatial navigation is reliant upon multiple sources of sensory information, fMRI spatial navigation paradigms only allow for visual input signals. Active walking as part of ecological study designs would provide proprioceptive and self-motion feedback signals as well as an improved field of view to participants. Such studies are necessary to complement the present findings. Previous research has indeed shown that navigation performance in older subjects is tightly coupled to the availability of multiple sources of sensory information (Adamo et al., 2012). Finally, future studies should take into consideration the role of sex and should include an intermediate age group in order to gain a finer understanding of the neural dynamics subtending spatial navigation across the lifespan (Grön et al., 2000; van der Ham and Claessen, 2020).

## 5. Conclusion

To conclude, the present study shed light on the possibility that navigational deficits in old age are linked to functional differences in brain areas involved in visual processing and to impaired representations of landmarks in temporal regions. This work helps towards a better comprehension of the neural dynamics subtending landmark-based navigation and it provides new insights on the impact of age-related spatial processing changes on navigation capabilities. We argue that approaching the study of spatial navigation in healthy and pathological aging from the perspective of visuo-perceptual abilities is a critical next step in the field. Neuroimaging methods coupled with virtual reality paradigms open up promising avenues to investigate age-related changes in navigation ability and to evaluate the benefits of training programs on older adults’ spatial autonomy and mobility.

## Author Contributions

Study design: SR, MB, CH, AA; Data acquisition: SR, MD; Data processing: SR, MD; Manuscript writing: SR, MD, MB, AA.

## Acknowledgments

The authors would like to express their gratitude to the men and women who took part in this study. We thank the Quinze-Vingts Hospital for allowing us to acquire the MRI data. We thank Konogan Baranton and Isabelle Poulain (*Essilor*) for the manufacturing of MRI-compatible glasses.

## Footnotes

1 All older participants scored 28 or above on the MMSE except one participant who scored 24. We decided to include this subject nonetheless as his extended neuropsychological evaluation was normal and no significant changes were detected when removing him from the fMRI analyses

## Bibliography

Adamo, D. E., Brice??o, E. M., Sindone, J. A., Alexander, N. B., and Moffat, S. D. (2012). Age differences in virtual environment and real world path integration. Front. Aging Neurosci. 4, 1–9. doi:10.3389/fnagi.2012.00026.

Allen, G. L., Kirasic, K. C., Dobson, S. H., Long, R. G., and Beck, S. (1996). Predicting environmental learning from spatial abilities: An indirect route. Intelligence 22, 327–355. doi:https://doi.org/10.1016/S0160-2896(96)90026-4.

Allison, S. L., Fagan, A. M., Morris, J. C., and Head, D. (2016). Spatial Navigation in Preclinical Alzheimer’s Disease. J. Alzheimers. Dis. 52, 77–90. doi:10.3233/JAD-150855.

Antonova, E., Parslow, D., Brammer, M., Dawson, G. R., Jackson, S. H. D., and Morris, R. G. (2009). Age-related neural activity during allocentric spatial memory. Memory 17, 125–143. doi:10.1080/09658210802077348.

Auger, S. D., and Maguire, E. A. (2018). Retrosplenial Cortex Indexes Stability beyond the Spatial Domain. J. Neurosci. 38, 1472–1481. doi:10.1523/JNEUROSCI.2602-17.2017.

Auger, S. D., Mullally, S. L., and Maguire, E. A. (2012). Retrosplenial cortex codes for permanent landmarks. PLoS One 7. doi:10.1371/journal.pone.0043620.

Auger, S. D., Zeidman, P., and Maguire, E. A. (2015). A central role for the retrosplenial cortex in de novo environmental learning. Elife 4, 1–26. doi:10.7554/eLife.09031.

Bécu, M., Sheynikhovich, D., Tatur, G., Agathos, C. P., Bologna, L. L., Sahel, J.-A., et al. (2020). Age-related preference for geometric spatial cues during real-world navigation. Nat. Hum. Behav. 4, 88–99. doi:10.1038/s41562-019-0718-z.

Bo, J., Peltier, S. J., Noll, D. C., and Seidler, R. D. (2011). Age differences in symbolic representations of motor sequence learning. Neurosci. Lett. 504, 68–72. doi:10.1016/j.neulet.2011.08.060.

Bonner, M. F., and Epstein, R. A. (2017). Coding of navigational affordances in the human visual system. Proc. Natl. Acad. Sci. U. S. A. 114, 4793–4798. doi:10.1073/pnas.1618228114.

Brewer, A. a., and Barton, B. (2012). Effects of healthy aging on human primary visual cortex. Health (Irvine. Calif). 04, 695–702. doi:10.4236/health.2012.429109.

Burke, S. N., Gaynor, L. S., Barnes, C. A., Bauer, R. M., Bizon, J. L., Roberson, E. D., et al. (2018). Shared Functions of Perirhinal and Parahippocampal Cortices: Implications for Cognitive Aging. Trends Neurosci. 41, 349–359. doi:10.1016/J.TINS.2018.03.001.

Caffo, A. O., Lopez, A., Spano, G., Serino, S., Cipresso, P., Stasolla, F., et al. (2018). Spatial reorientation decline in aging: the combination of geometry and landmarks. Aging Ment. Health 22, 1372–1383. doi:10.1080/13607863.2017.1354973.

Cheng, K., and Newcombe, N. S. (2005). Is there a geometric module for spatial orientation? Squaring theory and evidence. Psychon. Bull. Rev. 12, 1–23. doi:10.3758/BF03196346.

Chersi, F., and Burgess, N. (2015). The Cognitive Architecture of Spatial Navigation: Hippocampal and Striatal Contributions. Neuron 88, 64–77. doi:10.1016/j.neuron.2015.09.021.

Chrastil, E. R. (2013). Neural evidence supports a novel framework for spatial navigation. Psychon. Bull. Rev. 20, 208–27. doi:10.3758/s13423-012-0351-6.

Ciaramelli, E., Rosenbaum, R. S., Solcz, S., Levine, B., and Moscovitch, M. (2010). Mental space travel: damage to posterior parietal cortex prevents egocentric navigation and reexperiencing of remote spatial memories. J. Exp. Psychol. Learn. Mem. Cogn. 36, 619–634. doi:10.1037/a0019181.

Clancy Dollinger, S. M. (1995). Mental rotation performance: Age, sex, and visual field differences. Dev. Neuropsychol. 11, 215–222. doi:10.1080/87565649509540614.

Colombo, D., Serino, S., Tuena, C., Pedroli, E., Dakanalis, A., Cipresso, P., et al. (2017). Egocentric and allocentric spatial reference frames in aging: A systematic review. Neurosci. Biobehav. Rev. 80, 605–621. doi:10.1016/j.neubiorev.2017.07.012.

Cona, G., and Scarpazza, C. (2019). Where is the “where” in the brain? A meta-analysis of neuroimaging studies on spatial cognition. Hum. Brain Mapp. 40, 1867–1886. doi:10.1002/hbm.24496.

Corsi, P. M. (1973). Human memory and the medial temporal region of the brain. 34.

Coughlan, G., Laczo, J., Hort, J., Minihane, A.-M., and Hornberger, M. (2018). Spatial navigation deficits - overlooked cognitive marker for preclinical Alzheimer disease? Nat. Rev. Neurol. 14, 496–506. doi:10.1038/s41582-018-0031-x.

Dillen, K. N. H., Jacobs, H. I. L., Kukolja, J., von Reutern, B., Richter, N., Onur, Ö. A., et al. (2016). Aberrant functional connectivity differentiates retrosplenial cortex from posterior cingulate cortex in prodromal Alzheimer’s disease. Neurobiol. Aging 44, 114–126. doi:https://doi.org/10.1016/j.neurobiolaging.2016.04.010.

Ekstrom, A. D. (2015). Why vision is important to how we navigate. Hippocampus 25, 731–735. doi:10.1002/hipo.22449.

Epstein, R. A. (2008). Parahippocampal and retrosplenial contributions to human spatial navigation. Trends Cogn. Sci. 12, 388–396. doi:10.1016/j.tics.2008.07.004.

Epstein, R. A., Patai, E. Z., Julian, J. B., and Spiers, H. J. (2017). The cognitive map in humans: spatial navigation and beyond. Nat. Neurosci. 20, 1504–1513. doi:10.1038/nn.4656.

Epstein, R. a, and Vass, L. K. (2014). Neural systems for landmark-based wayfinding in humans. Philos. Trans. R. Soc. Lond. B. Biol. Sci. 369, 20120533. doi:10.1098/rstb.2012.0533.

Fjell, A. M., McEvoy, L., Holland, D., Dale, A. M., and Walhovd, K. B. (2014). What is normal in normal aging? Effects of aging, amyloid and Alzheimer’s disease on the cerebral cortex and the hippocampus. Prog. Neurobiol. 117, 20–40. doi:10.1016/j.pneurobio.2014.02.004.

Folstein, M. F., Folstein, S. E., and McHugh, P. R. (1975). “Mini-mental state”. A practical method for grading the cognitive state of patients for the clinician. J. Psychiatr. Res. 12, 189–198.

Foo, P., Warren, W. H., Duchon, A., and Tarr, M. J. (2005). Do humans integrate routes into a cognitive map? Map-versus landmark-based navigation of novel shortcuts. J. Exp. Psychol. Learn. Mem. Cogn. 31, 195–215. doi:10.1037/0278-7393.31.2.195.

Foster, T. C., Defazio, R. A., and Bizon, J. L. (2012). Characterizing cognitive aging of spatial and contextual memory in animal models. Front. Aging Neurosci. 4, 12. doi:10.3389/fnagi.2012.00012.

Friston, K. J., Holmes, A. P., Worsley, K. J., Poline, J. P., Frith, C. D., and Frackowiak, R. S. J. (1995). Statistical parametric maps in functional imaging: a general linear approach. Hum. Brain Mapp. 2, 189–210.

Gazova, I., Laczó, J., Rubinova, E., Mokrisova, I., Hyncicova, E., Andel, R., et al. (2013). Spatial navigation in young versus older adults. Front. Aging Neurosci. 5, 94. doi:10.3389/fnagi.2013.00094.

Gazova, I., Vlcek, K., Laczó, J., Nedelska, Z., Hyncicova, E., Mokrisova, I., et al. (2012). Spatial navigation—a unique window into physiological and pathological aging. Front. Aging Neurosci. 4, 16. doi:10.3389/fnagi.2012.00016.

Giocomo, L. M. (2016). Environmental boundaries as a mechanism for correcting and anchoring spatial maps. J. Physiol. 594, 6501–6511. doi:10.1113/JP270624.

Glasser, M. F., Sotiropoulos, S. N., Wilson, J. A., Coalson, T. S., Fischl, B., Andersson, J. L., et al. (2013). The minimal preprocessing pipelines for the Human Connectome Project. Neuroimage 80, 105–124. doi:10.1016/j.neuroimage.2013.04.127.

Goodroe, S. C., Starnes, J., and Brown, T. I. (2018). The Complex Nature of Hippocampal-Striatal Interactions in Spatial Navigation. Front. Hum. Neurosci. 12, 250. doi:10.3389/fnhum.2018.00250.

Greene, N. R., and Naveh-Benjamin, M. (2020). A Specificity Principle of Memory: Evidence From Aging and Associative Memory. Psychol. Sci., 0956797620901760. doi:10.1177/0956797620901760.

Grön, G., Wunderlich, a P., Spitzer, M., Tomczak, R., and Riepe, M. W. (2000). Brain activation during human navigation: gender-different neural networks as substrate of performance. Nat. Neurosci. 3, 404–8. doi:10.1038/73980.

Harris, M. A., Wiener, J. M., and Wolbers, T. (2012). Aging specifically impairs switching to an allocentric navigational strategy. Front. Aging Neurosci. 4, 1–9. doi:10.3389/fnagi.2012.00029.

Harris, M. A., and Wolbers, T. (2012). Ageing effects on path integration and landmark navigation. Hippocampus 22, 1770–1780. doi:10.1002/hipo.22011.

Hartmeyer, S., Grzeschik, R., Wolbers, T., and Wiener, J. M. (2017). The Effects of Attentional Engagement on Route Learning Performance in a Virtual Environment: An Aging Study. Front. Aging Neurosci. 9, 235. Available at: https://www.frontiersin.org/article/10.3389/fnagi.2017.00235.

Herweg, N. A., and Kahana, M. J. (2018). Spatial Representations in the Human Brain. Front. Hum. Neurosci. 12, 297. doi:10.3389/fnhum.2018.00297.

Hirsch, P., Schwarzkopp, T., Declerck, M., Reese, S., and Koch, I. (2016). Age-related differences in task switching and task preparation: Exploring the role of task-set competition. Acta Psychol. (Amst). doi:10.1016/j.actpsy.2016.06.008.

Iachini, T., Poderico, C., Ruggiero, G., and Iavarone, A. (2005). Age differences in mental scanning of locomotor maps. Disabil. Rehabil. 27, 741–752. doi:10.1080/09638280400014782.

Iaria, G., Palermo, L., Committeri, G., and Barton, J. J. S. (2009). Age differences in the formation and use of cognitive maps. Behav. Brain Res. 196, 187–191. doi:10.1016/j.bbr.2008.08.040.

Iaria, G., Petrides, M., Dagher, A., Pike, B., and Bohbot, V. D. (2003). Cognitive strategies dependent on the hippocampus and caudate nucleus in human navigation: variability and change with practice. J. Neurosci. 23, 5945–5952. doi:23/13/5945 [pii].

Igloi, K., Doeller, C. F., Berthoz, A., Rondi-Reig, L., and Burgess, N. (2010). Lateralized human hippocampal activity predicts navigation based on sequence or place memory. Proc. Natl. Acad. Sci. U. S. A. 107, 14466–14471. doi:10.1073/pnas.1004243107/-/DCSupplemental.www.pnas.org/cgi/doi/10.1073/pnas.1004243107.

Igloi, K., Doeller, C. F., Paradis, A.-L., Benchenane, K., Berthoz, A., Burgess, N., et al. (2015). Interaction Between Hippocampus and Cerebellum Crus I in Sequence-Based but not Place-Based Navigation. Cereb. Cortex 25, 4146–4154. doi:10.1093/cercor/bhu132.

Igloi, K., Doeller, C. F., Paradis, A.-L., Benchenane, K., Berthoz, A., Burgess, N., et al. (2014). Interaction Between Hippocampus and Cerebellum Crus I in Sequence-Based but not Place-Based Navigation. Cereb. Cortex, bhu132-. doi:10.1093/cercor/bhu132.

Ito, H. T. (2018). Prefrontal-hippocampal interactions for spatial navigation. Neurosci. Res. 129, 2–7. doi:10.1016/j.neures.2017.04.016.

Janzen, G., and van Turennout, M. (2004). Selective neural representation of objects relevant for navigation. Nat. Neurosci. 7, 673–677. doi:10.1038/nn1257.

Javadi, A.-H., Emo, B., Howard, L. R., Zisch, F. E., Yu, Y., Knight, R., et al. (2017). Hippocampal and prefrontal processing of network topology to simulate the future. Nat. Commun. 8, 14652. doi:10.1038/ncomms14652.

Julian, J. B., Keinath, A. T., Marchette, S. A., and Epstein, R. A. (2018). The Neurocognitive Basis of Spatial Reorientation. Curr. Biol. 28, R1059–R1073. doi:10.1016/j.cub.2018.04.057.

Julian, J. B., Ryan, J., Hamilton, R. H., Epstein, R. A., Julian, J. B., Ryan, J., et al. (2016). The Occipital Place Area Is Causally Involved in Representing Environmental Boundaries during Navigation. Curr. Biol., 1–6. doi:10.1016/j.cub.2016.02.066.

Kamps, F. S., Julian, J. B., Kubilius, J., Kanwisher, N., and Dilks, D. D. (2016). The occipital place area represents the local elements of scenes. Neuroimage 132, 417–424. doi:10.1016/j.neuroimage.2016.02.062.

Kauffmann, L., Ramanoël, S., and Peyrin, C. (2014). The neural bases of spatial frequency processing during scene perception. Front. Integr. Neurosci. 8. doi:10.3389/fnint.2014.00037.

Kimura, K., Reichert, J. F., Kelly, D. M., and Moussavi, Z. (2019). Older Adults Show Less Flexible Spatial Cue Use When Navigating in a Virtual Reality Environment Compared With Younger Adults. Neurosci. Insights 14, 2633105519896803. doi:10.1177/2633105519896803.

Kirasic, K. C. (1991). Spatial cognition and behavior in young and elderly adults: implications for learning new environments. Psychol. Aging 6, 10–18. doi:10.1037//0882-7974.6.1.10.

Konishi, K., Etchamendy, N., Roy, S., Marighetto, A., Rajah, N., and Bohbot, V. D. (2013). Decreased functional magnetic resonance imaging activity in the hippocampus in favor of the caudate nucleus in older adults tested in a virtual navigation task. Hippocampus 23, 1005–1014. doi:10.1002/hipo.22181.

Kozhevnikov, M., and Hegarty, M. (2001). A dissociation between object manipulation spatial ability and spatial orientation ability. Mem. Cognit. 29, 745–756. doi:10.3758/bf03200477.

Kozhevnikov, M., Motes, M. A., Rasch, B., and Blajenkova, O. (2006). Perspective-taking vs. mental rotation transformations and how they predict spatial navigation performance. Appl. Cogn. Psychol. 20, 397–417. doi:10.1002/acp.1192.

Kravitz, D. J., Saleem, K. S., Baker, C. I., Ungerleider, L. G., and Mishkin, M. (2013). The ventral visual pathway: an expanded neural framework for the processing of object quality. Trends Cogn. Sci. 17, 26–49. doi:https://doi.org/10.1016/j.tics.2012.10.011.

Kuhn, S., and Gallinat, J. (2014). Segregating cognitive functions within hippocampal formation: a quantitative meta-analysis on spatial navigation and episodic memory. Hum. Brain Mapp. 35, 1129–1142. doi:10.1002/hbm.22239.

Laczo, J., Andel, R., Nedelska, Z., Vyhnalek, M., Vlcek, K., Crutch, S., et al. (2017). Exploring the contribution of spatial navigation to cognitive functioning in older adults. Neurobiol. Aging 51, 67–70. doi:10.1016/j.neurobiolaging.2016.12.003.

Laczo, J., Parizkova, M., and Moffat, S. D. (2018). Spatial navigation, aging and Alzheimer’s disease. Aging (Albany. NY). 10, 3050–3051. doi:10.18632/aging.101634.

Lester, A. W., Moffat, S. D., Wiener, J. M., Barnes, C. A., and Wolbers, T. (2017). The Aging Navigational System. Neuron 95, 1019–1035. doi:10.1016/j.neuron.2017.06.037.

Li, A. W. Y., and King, J. (2019). Spatial memory and navigation in ageing: A systematic review of MRI and fMRI studies in healthy participants. Neurosci. Biobehav. Rev. 103, 33–49. doi:10.1016/j.neubiorev.2019.05.005.

Lithfous, S., Dufour, A., and Despres, O. (2013). Spatial navigation in normal aging and the prodromal stage of Alzheimer’s disease: insights from imaging and behavioral studies. Ageing Res. Rev. 12, 201–213. doi:10.1016/j.arr.2012.04.007.

Litman, L., Awipi, T., and Davachi, L. (2009). Category-specificity in the human medial temporal lobe cortex. Hippocampus 19, 308–319. doi:10.1002/hipo.20515.

Marchette, S. a, Vass, L. K., Ryan, J., and Epstein, R. a (2015). Outside Looking In: Landmark Generalization in the Human Navigational System. J. Neurosci. 35, 14896–908. doi:10.1523/JNEUROSCI.2270-15.2015.

Mazaika, P.; Hoeft, F.; Glover, GH.; Reiss, A.. (2009). Methods and Software for fMRI analysis for Clinical Subjects. in Organization for Human Brain Mapping (San Francisco, CA, USA).

Meneghetti, C., Muffato, V., Borella, E., and De Beni, R. (2018). Map Learning in Normal Aging: The Role of Individual Visuo-Spatial Abilities and Implications. Curr. Alzheimer Res. 15, 205–218. doi:10.2174/1567205014666171030113515.

Merhav, M., Riemer, M., and Wolbers, T. (2019). Spatial updating deficits in human aging are associated with traces of former memory representations. Neurobiol. Aging 76, 53–61. doi:10.1016/j.neurobiolaging.2018.12.010.

Merhav, M., and Wolbers, T. (2019). Aging and spatial cues influence the updating of navigational memories. Sci. Rep. 9, 11469. doi:10.1038/s41598-019-47971-2.

Meulenbroek, O., Petersson, K. M., Voermans, N., Weber, B., and Fernandez, G. (2004). Age differences in neural correlates of route encoding and route recognition. Neuroimage 22, 1503–1514. doi:10.1016/j.neuroimage.2004.04.007.

Miniaci, M. C., and De Leonibus, E. (2018). Missing the egocentric spatial reference: a blank on the map. F1000Research 7, 168. doi:10.12688/f1000research.13675.1.

Mitchell, A. S., Czajkowski, R., Zhang, N., Jeffery, K., and Nelson, A. J. D. (2018). Retrosplenial cortex and its role in spatial cognition. Brain Neurosci. Adv. 2, 2398212818757098. doi:10.1177/2398212818757098.

Moffat, S. D. (2009). Aging and spatial navigation: What do we know and where do we go? Neuropsychol. Rev. 19, 478–489. doi:10.1007/s11065-009-9120-3.

Moffat, S. D., Elkins, W., and Resnick, S. M. (2006). Age differences in the neural systems supporting human allocentric spatial navigation. Neurobiol. Aging 27, 965–972. doi:10.1016/j.neurobiolaging.2005.05.011.

Moffat, S. D., and Resnick, S. M. (2002). Effects of age on virtual environment place navigation and allocentric cognitive mapping. Behav. Neurosci. 116, 851–859. doi:10.1037/0735-7044.116.5.851.

Muffato, V., Meneghetti, C., and De Beni, R. (2020). The role of visuo-spatial abilities in environment learning from maps and navigation over the adult lifespan. Br. J. Psychol. 111, 70–91. doi:10.1111/bjop.12384.

Nau, M., Julian, J. B., and Doeller, C. F. (2018). How the Brain’s Navigation System Shapes Our Visual Experience. Trends Cogn. Sci. 22, 810–825. doi:10.1016/j.tics.2018.06.008.

Ohta, R. J., Walsh, D. A., and Krauss, I. K. (1981). Spatial perspective-taking ability in young and elderly adults. Exp. Aging Res. 7, 45–63. doi:10.1080/03610738108259785.

Packard, M. G., and Goodman, J. (2013). Factors that influence the relative use of multiple memory systems. Hippocampus 23, 1044–1052. doi:10.1002/hipo.22178.

Perrochon, A., Mandigout, S., Petruzzellis, S., Soria Garcia, N., Zaoui, M., Berthoz, A., et al. (2018). The influence of age in women in visuo-spatial memory in reaching and navigation tasks with and without landmarks. Neurosci. Lett. 684, 13–17. doi:10.1016/j.neulet.2018.06.054.

Persichetti, A. S., and Dilks, D. D. (2016). Perceived egocentric distance sensitivity and invariance across scene-selective cortex. Cortex. 77, 155–163. doi:10.1016/j.cortex.2016.02.006.

Persichetti, A. S., and Dilks, D. D. (2018). Dissociable Neural Systems for Recognizing Places and Navigating through Them. J. Neurosci. 38, 10295 LP – 10304. doi:10.1523/JNEUROSCI.1200-18.2018.

Persichetti, A. S., and Dilks, D. D. (2019). Distinct representations of spatial and categorical relationships across human scene-selective cortex. Proc. Natl. Acad. Sci. 116, 21312 LP 21317. doi:10.1073/pnas.1903057116.

Picucci, L., Caffò, A. O., and Bosco, A. (2009). Age and sex differences in a virtual version of the reorientation task. Cogn. Process. doi:10.1007/s10339-009-0321-8.

Ramanoël, S., Kauffmann, L., Cousin, E., Dojat, M., and Peyrin, C. (2015). Age-related differences in spatial frequency processing during scene categorization. PLoS One. doi:10.1371/journal.pone.0134554.

Ramanoël, S., York, E., Le Petit, M., Lagrene, K., Habas, C., and Arleo, A. (2019). Age-Related Differences in Functional and Structural Connectivity in the Spatial Navigation Brain Network. Front. Neural Circuits 13, 69. doi:10.3389/fncir.2019.00069.

Ratliff, K. R., and Newcombe, N. S. (2008). Reorienting when cues conflict: Evidence for an adaptive-combination view. Psychol. Sci. doi:10.1111/j.1467-9280.2008.02239.x.

Rochefort, C., Lefort, J. M., and Rondi-Reig, L. (2013). The cerebellum: a new key structure in the navigation system. Front. Neural Circuits 7, 35. doi:10.3389/fncir.2013.00035.

Rodgers, M. K., Sindone, J. A., and Moffat, S. D. (2012). Effects of age on navigation strategy. Neurobiol. Aging 33, 202.e15–202.e22. doi:10.1016/j.neurobiolaging.2010.07.021.

Sack, A. T. (2009). Parietal cortex and spatial cognition. Behav. Brain Res. 202, 153–161. doi:https://doi.org/10.1016/j.bbr.2009.03.012.

Schuck, N. W., Doeller, C. F., Polk, T. A., Lindenberger, U., and Li, S.-C. (2015a). Human aging alters the neural computation and representation of space. Neuroimage I, 141–150. doi:10.1016/j.neuroimage.2015.05.031.

Schuck, N. W., Doeller, C. F., Polk, T. A., Lindenberger, U., and Li, S. C. (2015b). Human aging alters the neural computation and representation of space. Neuroimage. doi:10.1016/j.neuroimage.2015.05.031.

Seghier, M. L. (2012). The Angular Gyrus. Neurosci. 19, 43–61. doi:10.1177/1073858412440596.

Sherrill, K. R., Chrastil, E. R., Ross, R. S., Erdem, U. M., Hasselmo, M. E., and Stern, C. E. (2015). Functional connections between optic flow areas and navigationally responsive brain regions during goal-directed navigation. Neuroimage 118, 386–396. doi:10.1016/j.neuroimage.2015.06.009.

Spiers, H. J., and Barry, C. (2015a). Neural systems supporting navigation. Curr. Opin. Behav. Sci. 1, 47–55. doi:10.1016/j.cobeha.2014.08.005.

Spiers, H. J., and Barry, C. (2015b). Neural systems supporting navigation. Curr. Opin. Behav. Sci. 1, 47–55. doi:https://doi.org/10.1016/j.cobeha.2014.08.005.

Stankiewicz, B. J., and Kalia, A. A. (2007). Acquisition of Structural Versus Object Landmark Knowledge. J. Exp. Psychol. Hum. Percept. Perform. doi:10.1037/0096-1523.33.2.378.

Sturz, B. R., Kilday, Z. A., and Bodily, K. D. (2013). Does constraining field of view prevent extraction of geometric cues for humans during virtual-environment reorientation? J. Exp. Psychol. Anim. Behav. Process. 39, 390–396. doi:10.1037/a0032543.

Sutton, J. E., Joanisse, M. F., and Newcombe, N. S. (2010). Spinning in the scanner: Neural correlates of virtual reorientation. J. Exp. Psychol. Learn. Mem. Cogn. 36, 1097–1107. doi:10.1037/a0019938.

Techentin, C., Voyer, D., and Voyer, S. D. (2014). Spatial Abilities and Aging: A Meta-Analysis. Exp. Aging Res. 40, 395–425. doi:10.1080/0361073X.2014.926773.

Tommasi, L., Chiandetti, C., Pecchia, T., Sovrano, V. A., and Vallortigara, G. (2012). From natural geometry to spatial cognition. Neurosci. Biobehav. Rev. 36, 799–824. doi:10.1016/j.neubiorev.2011.12.007.

United Nations, Departement of Economic and Social Affairs, Population Division (2019). World population ageing. Highlight (ST/ESA/SER.A/430).

van der Ham, I. J. M., Baalbergen, H., van der Heijden, P. G. M., Postma, A., Braspenning, M., and van der Kuil, M. N. A. (2015). Distance comparisons in virtual reality: effects of path, context, and age. Front. Psychol. 6, 1103. doi:10.3389/fpsyg.2015.01103.

van der Ham, I. J. M., and Claessen, M. H. G. (2020). How age relates to spatial navigation performance: Functional and methodological considerations. Ageing Res. Rev. 58, 101020. doi:https://doi.org/10.1016/j.arr.2020.101020.

Vandenberg, S. G., and Kuse, A. R. (1978). Mental rotations, a group test of three-dimensional spatial visualization. Percept. Mot. Skills 47, 599–604. doi:10.2466/pms.1978.47.2.599.

Vann, S. D., Aggleton, J. P., and Maguire, E. A. (2009). What does the retrosplenial cortex do? Nat. Rev. Neurosci. 10, 792–802. doi:10.1038/nrn2733.

Wandell, B. A., Brewer, A. A., and Dougherty, R. F. (2005). Visual field map clusters in human cortex. Philos. Trans. R. Soc. Lond. B. Biol. Sci. 360, 693–707. doi:10.1098/rstb.2005.1628.

Weiskopf, N., Hutton, C., Josephs, O., and Deichmann, R. (2006). Optimal EPI parameters for reduction of susceptibility-induced BOLD sensitivity losses: A whole-brain analysis at 3 T and 1.5 T. Neuroimage. doi:10.1016/j.neuroimage.2006.07.029.

Wiener, J. M., de Condappa, O., Harris, M. A., and Wolbers, T. (2013). Maladaptive Bias for Extrahippocampal Navigation Strategies in Aging Humans. J. Neurosci. doi:10.1523/jneurosci.0717-12.2013.

Wiener, J. M., Kmecova, H., and de Condappa, O. (2012). Route repetition and route retracing: Effects of cognitive aging. Front. Aging Neurosci. 4, 1–7. doi:10.3389/fnagi.2012.00007.

Wilkniss, S. M., Jones, M. G., Korol, D. L., Gold, P. E., and Manning, C. A. (1997). Age-related differences in an ecologically based study of route learning. Psychol. Aging 12, 372–375. doi:10.1037/0882-7974.12.2.372.

Wolbers, T., and Büchel, C. (2005). Dissociable Retrosplenial and Hippocampal Contributions to Successful Formation of Survey Representations. J. Neurosci. 25, 3333–3340. doi:10.1523/JNEUROSCI.4705-04.2005.

Wolbers, T., and Hegarty, M. (2010). What determines our navigational abilities? Trends Cogn. Sci. 14, 138–146. doi:10.1016/j.tics.2010.01.001.

Zajac, L., Burte, H., Taylor, H. A., and Killiany, R. (2019). Self-reported navigation ability is associated with optic flow-sensitive regions’ functional connectivity patterns during visual path integration. Brain Behav. 9, e01236. doi:10.1002/brb3.1236.

Zhong, J. Y., and Moffat, S. D. (2016). Age-Related Differences in Associative Learning of Landmarks and Heading Directions in a Virtual Navigation Task. Front. Aging Neurosci. 8, 122. doi:10.3389/fnagi.2016.00122.

Zhong, J. Y., and Moffat, S. D. (2018). Extrahippocampal Contributions to Age-Related Changes in Spatial Navigation Ability. Front. Hum. Neurosci. 12, 272. doi:10.3389/fnhum.2018.00272.

